# Differential viral RNA methylation contributes to pathogen blocking in *Wolbachia*-colonized arthropods

**DOI:** 10.1101/2021.03.26.437201

**Authors:** Tamanash Bhattacharya, Liewei Yan, Hani Zaher, Irene L.G. Newton, Richard W. Hardy

## Abstract

Arthropod endosymbiont *Wolbachia pipientis* is part of a global biocontrol strategy to reduce the replication of mosquito-borne RNA viruses such as alphaviruses. We previously demonstrated the importance of a host cytosine methyltransferase, DNMT2, in *Drosophila* and viral RNA as a cellular target during pathogen-blocking. Here we report on the role of DNMT2 in *Wolbachia-*induced alphavirus inhibition in *Aedes* species. Expression of DNMT2 in mosquito tissues, including the salivary glands, is elevated upon virus infection. Notably, this is suppressed in *Wolbachia-colonized* animals, coincident with reduced virus replication and decreased infectivity of progeny virus. Ectopic expression of DNMT2 in cultured *Aedes* cells is proviral, increasing progeny virus infectivity, and this effect of DNMT2 on virus replication and infectivity is dependent on its methyltransferase activity. Finally, examining the effects of *Wolbachia* on modifications of viral RNA by LC-MS show a decrease in the amount of 5-methylcytosine modification consistent with the down-regulation of DNMT2 in *Wolbachia* colonized mosquito cells and animals. Collectively, our findings support the conclusion that disruption of 5-methylcytosine modification of viral RNA is a vital mechanism operative in pathogen blocking. These data also emphasize the essential role of epitranscriptomic modifications in regulating fundamental alphavirus replication and transmission processes.

## Introduction

Viruses are remarkably adept at using a limited set of viral factors to replicate in vastly different host cell environments. This ability is vital for the success of zoonotic arboviruses, which encounter physiologically and ecologically distinct invertebrate and vertebrate hosts during transmission. As these viruses oscillate between vertebrate and arthropod hosts, the progeny virions reared in one host cell context are primed for the next, predicating successful transmission. However, arbovirus transmission events are influenced by many host-specific biotic and abiotic factors (2–6). Recent studies have identified the vector microbiome as a critical biotic factor influencing arbovirus transmission (5, 6). One notable member of this microbial population is the arthropod endosymbiont *Wolbachia pipientis*, which dramatically impacts the transmission of multiple zoonotic arboviruses, a phenomenon termed “pathogen-blocking” (PB) (7–15). *Wolbachia* is transmitted transovarially and induces a wide range of reproductive manipulations in its host (16, 17). For example, *Wolbachia’s* presence results in sperm-egg incompatibility between colonized and non-colonized individuals, mediated by a bacterially-encoded toxin-antitoxin system (17). This phenomenon, known as cytoplasmic incompatibility (CI), allows *Wolbachia* to be inherited at a rate higher than Mendelian inheritance, like a natural gene drive. Over the last decade, scientists have leveraged this property to deploy *Wolbachia* as a novel vector control agent with the aim of either suppressing or replacing the local mosquito population (18). Recent data suggest that *Wolbachia* release programs significantly reduce the transmission of Dengue virus (DENV) in endemic regions across 11 territories in Asia and Latin America (19, 20). Remarkably, however, despite its success, the underlying cellular mechanism of pathogen-blocking remains unidentified.

We recently showed that viral RNA is a cellular target of *Wolbachia-mediated* inhibition and that loss in progeny virus infectivity occurs at the level of the encapsidated virion RNA, which is compromised in its ability to replicate in naïve vertebrate cells (1). Mosquito-derived viruses, reared in the presence of *Wolbachia*, are less infectious when seeded in either mosquito or vertebrate cells. These results suggest a transgenerational mechanism by which *Wolbachia* limits virus dissemination within the mosquito and subsequent transmission into vertebrates (1). We, therefore, speculated that factor(s) regulating pathogen blocking likely target the viral plus sense RNA genome, a feature shared between all viruses susceptible to *Wolbachia-mediated* inhibition. Notably, the presence of *Wolbachia* reduces the infectivity of the encapsidated virion RNA in mammalian cells (1). This observation alone suggests one or more RNA-targeting factors may be responsible for compromising viral RNA replication in *Wolbachia-colonized* arthropod cells, as well as in mammalian cells, which are devoid of *Wolbachia*. Prior work has implicated mosquito exonuclease in pathogen-blocking, which is in line with the reduced half-life of incoming viral RNAs in *Wolbachia-colonized* cells (1, 21). While faster degradation of viral RNA explains the observed reduction in virus replication in arthropod cells, it does not explain reduced replication in mammalian cells. In light of these findings, we sought to focus our attention on the RNA cytosine methyltransferase DNMT2. Our prior study demonstrated that DNMT2 is essential for pathogen-blocking in fruit flies (9). As an RNA modifying protein, DNMT2 does not directly antagonize viral RNA replication but instead influences the cellular fate of its target(s). Both arthropods and mammals encode proteins capable of interpreting epitranscriptomic signatures on different RNA species, so we hypothesized that DNMT2-mediated modifications to the viral RNA in *Wolbachia-*colonized arthropods impact virus replication in arthropod and mammalian cells.

To test this hypothesis, we investigated whether DNMT2 is essential for *Wolbachia-mediated* pathogen blocking in mosquitoes. Additionally, we ask whether this MTase is functionally crucial to virus regulation in the absence of *Wolbachia*. Given DNMT2’s biological role as a cellular RNA cytosine methyltransferase, we further examined the possibility of m5C modification of viral RNA in mosquito cells and whether viral RNA is differentially modified in the presence and absence of *Wolbachia* in mosquito cells. We find that *Wolbachia* and viruses differentially influence MTase expression in mosquitoes. Specifically, the presence of the virus leads to elevated MTase expression, which is proviral in mosquito cells. In contrast, the presence of *Wolbachia* downregulates MTase levels to seemingly disrupt this proviral state, contributing to virus inhibition as well as reduced progeny virus infectivity. Furthermore, the proviral effect is dependent on the catalytic activity of DNMT2. Finally, as a consequence of this downregulation and DNMT2’s role as an RNA cytosine MTase, we show that the presence of *Wolbachia* in cells results in a reduced abundance of 5-methylcytosine (m5C) modification of progeny viral RNA. These changes imply that m5C modifications play a role in regulating viral RNA infectivity in mammalian cells. In summary, our findings highlight a previously underappreciated role of RNA methylation in alphavirus replication, with important implications for virus dissemination and transmission. Overall, our results indicate a role of the viral epitranscriptome as regulatory signatures capable of influencing the transmission of other arboviruses.

## Results

### Virus and *Wolbachia* differentially modulate *Aedes* DNMT2 expression

*Wolbachia* in *Aedes* mosquitoes is associated with reduced DNMT2 *(AMt2)* expression (22). We, therefore, examined the expression of *AMt2* in *w*AlbB-colonized *Aedes aegypti* mosquitoes. We chose to assess *in vivo AMt2* expression changes in whole mosquitos (which would give us a sense of the *Mt2* environment encountered by disseminating viruses) or in dissected salivary glands (the tissue important for transmission to the vertebrate host).

We measured *AMt2* expression in female *Aedes aegypti* mosquitoes colonized with and without *Wolbachia* (*w*AlbB) five days post eclosion, forty-eight hours following bloodmeals with and without Sindbis virus (SINV) (Fig 1B). The presence of both endosymbiont and virus was associated with altered *AMt2* expression (Two-way ANOVA, p < 0.0001), with a nearly 100-fold increase in *AMt2* levels in *Wolbachia-*free mosquitoes that received an infectious virus-containing blood meal (Fig 1B. W−/V− compared to W−/V+; Two-way ANOVA, p < 0.0001). In contrast, we found *Wolbachia* to reduce *AMt2* expression in mock-infected individuals (W+/V−) by approximately 5-fold (Fig 1B, W−/V− compared to W+/V−). Importantly, we also observed low *AMt2* expression in *Wolbachia-*colonized mosquitoes post-infectious (V+) bloodmeal, indicating that *Wolbachia* prevents virus-induced stimulation of *AMt2* expression and that these levels are maintained during infection. Virus replication in *Wolbachia-*colonized mosquitoes, therefore, occurs in a low *AMt2* environment (Fig 1B. W−/V− compared to W+/V+). This pattern of reduced *AMt2* expression was also observed *ex vivo* in cultured *Aedes albopictus-derived* mosquito cells colonized with both a native (*w*AlbB strain in Aa23 cells) and a non-native *Wolbachia* (*w*Mel in RML12 cells) strain (Fig S1).

**Fig 1.**
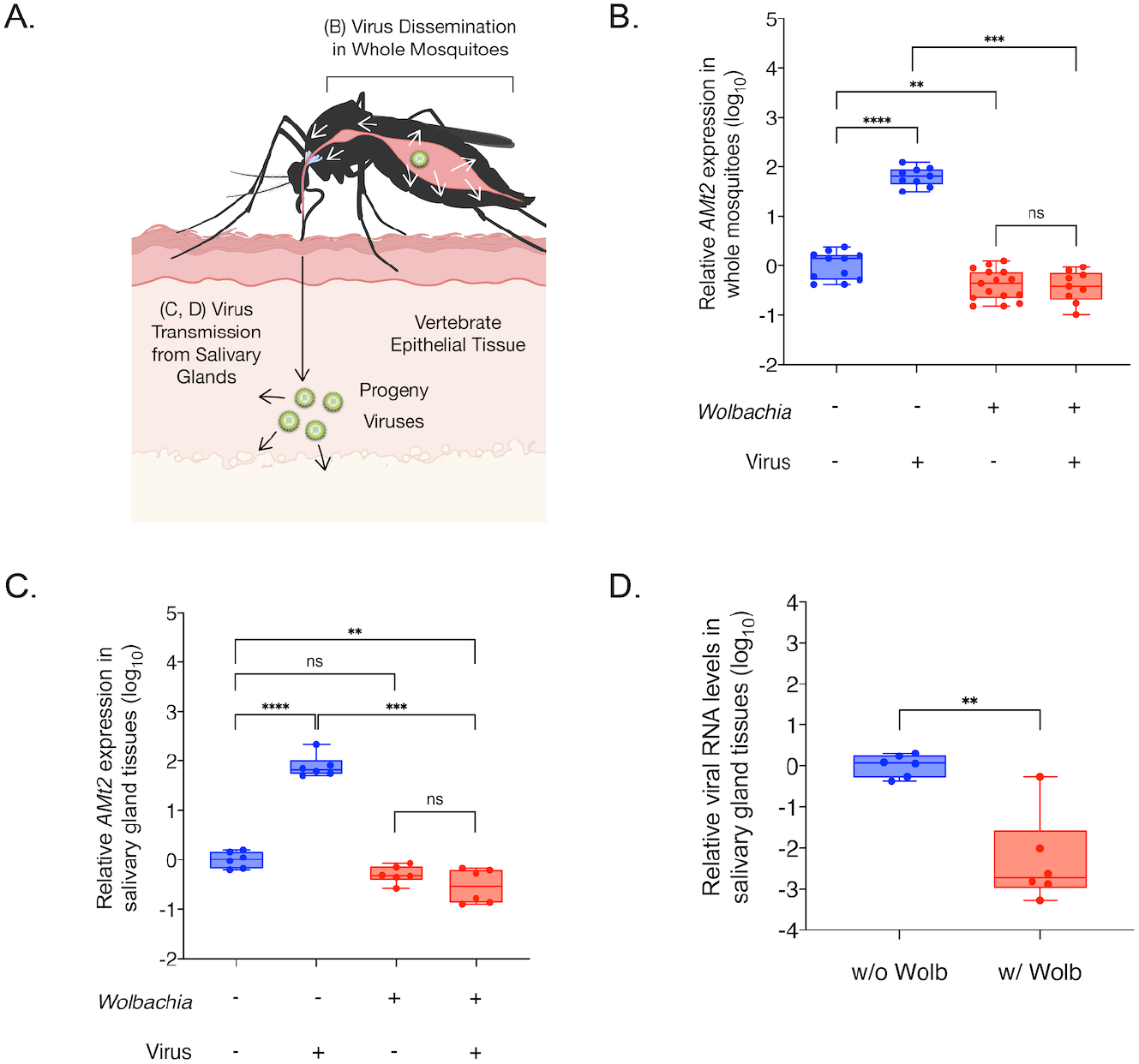
Virus and *Wolbachia* each differentially modulate expression of the RNA methyltransferase gene DNMT2 in mosquitoes. (A) Schematic of virus dissemination (white arrows) within mosquito tissues and transmission into vertebrate host (black arrow). (B) *AMt2* expression was measured 5 days post bloodmeal in whole female mosquitoes using qRT-PCR with and without SINV. Error bars represent standard error of mean (SEM) of biological replicates. Two-way ANOVA with Tukey’s post-hoc test of multivariate comparisons. (C) *AMt2* expression measured in dissected salivary gland tissues collected from female mosquitoes with and *Wolbachia-free* 5 days post bloodmeal with or without SINV. Error bars represent standard error of mean (SEM) of biological replicates. Two-way ANOVA with Tukey’s post-hoc test of multivariate comparisons. (D) Viral RNA levels were quantified in dissected salivary gland tissues with and *Wolbachia-free* using qRT-PCR at 5 days post infectious blood meal with SINV. Unpaired, student’s t-test, error bars represent standard error of mean (SEM) of biological replicates. For all panels: ****P < 0.0001; ***P < 0.001; **P < 0.01; ns = non-significant.

We next quantified *AMt2* expression in isolated salivary gland tissues from five-day-old *Aedes aegypti* mosquitoes colonized with or *Wolbachia-free* (*w*AlbB), post bloodmeal with (V+) or without (V−) SINV (Fig 1C). *AMt2* expression in the salivary gland tissues post-infectious bloodmeal was elevated nearly 100-fold, similar to the increase induced by the virus in whole mosquitoes (Fig 1C. W−/V− compared to W−/V+). As before, the presence of *Wolbachia* alone was associated with lower *AMt2* expression. However, this difference was not statistically significant (Fig 1C. W−/V− compared to W+/V−). Importantly, however, *Wolbachia* did prevent SINV-induced *AMt2* upregulation, reducing it 100-1000 fold (Fig 1C. W−/V+ compared to W+/V+). Under these conditions, we also observed a significant 2 to 3 log_10_ reduction in viral RNA in the salivary gland tissues (Fig 1D). It should be noted that our observations regarding the effect of *Wolbachia* (*w*AlbB) or SINV on *AMt2* expression are analogous to previous reports that describe differential *AMt2* expression in the presence of the flavivirus DENV-2 and *Wolbachia* (*w*Mel) in *Aedes aegypti* mosquitoes (22).

### DNMT2 promotes virus infection in mosquito cells

The positive correlation between *AMt2* expression and SINV genome replication in *Aedes* mosquitoes (Fig 1B-D) led us to examine whether there is a functional consequence of elevated MTase expression on virus infection in these insects. We therefore ectopically expressed *AMt2* and assessed its effect on virus infection in cultured *Aedes albopictus* cells (Fig 2A), using azacytidine-immunoprecipitation (AZA-IP) to determine whether viral RNA in the cell is a direct DNMT2 target. *Wolbachia-free Aedes albopictus* (C710) cells were transfected with an epitope-tagged *AMt2* expression vector (FLAG-*AMt2*) or control vector (FLAG-empty) for 48 hours before infection with SINV at an MOI of 10. After 24 hours post-infection, cells were labeled with a cytosine analog, 5-Azacytidine (5-AZAC), for 18 hours to incorporate the label into newly synthesized cellular and viral RNA. We reasoned that if mosquito DNMT2 directly targets viral RNAs for methylation, the presence of 5-AZAC in the RNA should covalently trap the enzyme forming a stable m5C-DNMT2-viral RNA complex, allowing co-immunoprecipitation of the RNA-protein complexes using anti-FLAG antibody (23). Targeted quantitative RT-PCR analyses of total immunoprecipitated RNA revealed enrichment of SINV RNA relative to a control host RNA transcript (GAPDH), confirming that viral RNA is indeed a direct MTase target in these cells (Fig 2B).

**Fig 2.**
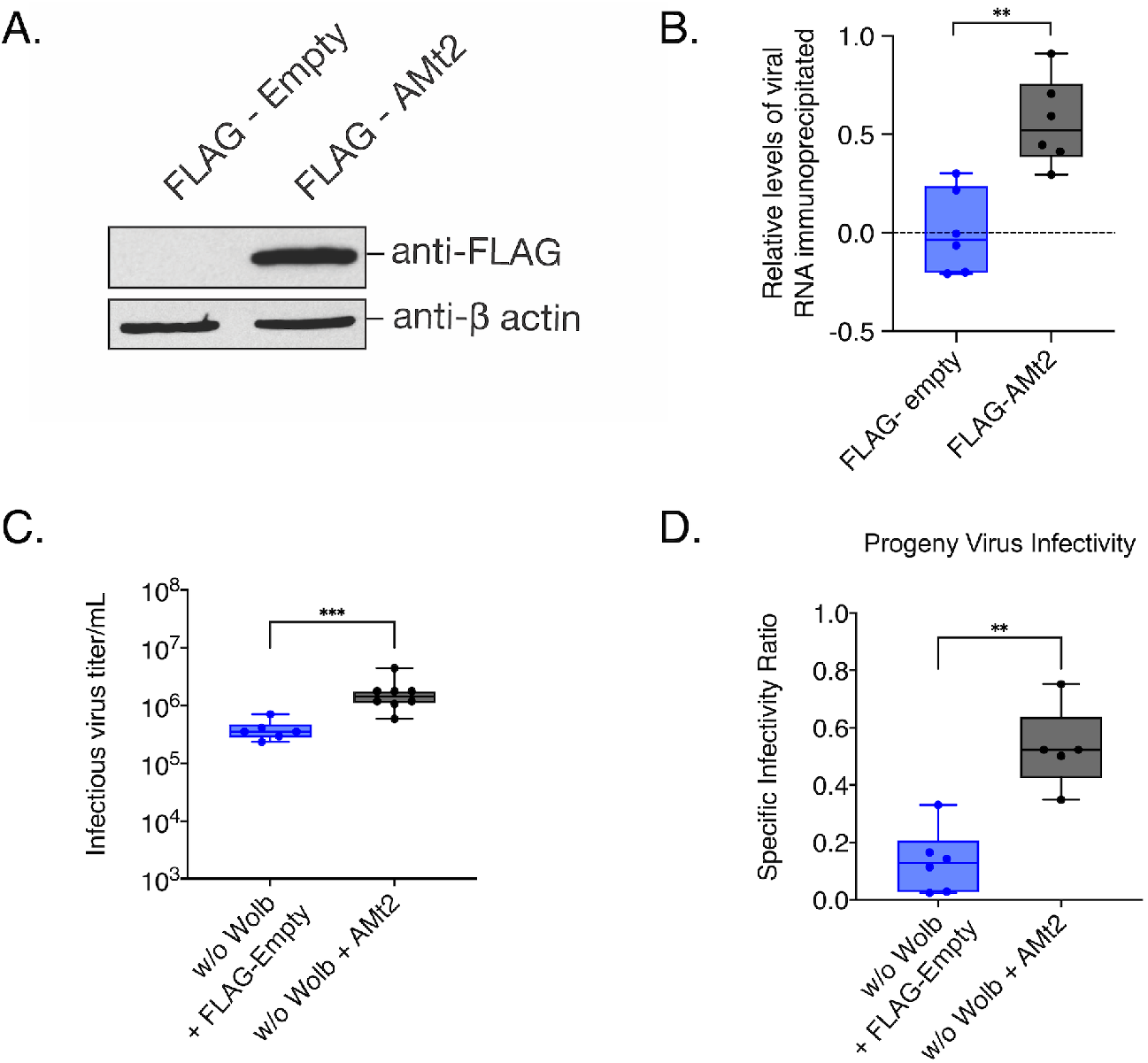
Overexpressing *AMt2* in mosquito cells improve progeny virus infectivity. (A) Western Blot of *Aedes albopictus* (C710) cells transfected with expression vector constructs with (FLAG-AMt2) or without (FLAG-empty) *AMt2*. Cytoplasmic lysates of cells were collected 48 hours post transfection and probed with anti-FLAG and anti-β actin antibodies. (B) Relative levels of SINV RNA recovered following AZA-IP of AMt2 in C710 cells was quantified using qRT-PCR. *Wolbachia-free* C710 mosquito cells were transfected with expression vectors FLAG-empty or FLAG-AMt2 for 48 hours prior to infection with SINV at MOI of 10. Cells were treated for approximately 18h with 5 μM 5-Azacytidine to covalently trap AMt2 with its target cellular RNA prior to RNA immunoprecipitation using anti-FLAG antibody. The horizontal dotted line represents the threshold set at 1 (log_10_). Unpaired two-tailed t-test with Welch’s correction, p = 0.0004, t = 4.216, df = 20 (C) Infectious progeny (PFU/mL) SINV produced from mosquito cells *Wolbachia-free* expressing either FLAG-empty (w/o Wolb) or FLAG-AMt2 (w/o Wolb + AMt2). Cells were transfected 48 hours prior to infection with SINV at MOI of 10. Infectious progeny viruses collected from supernatants 48 hours post-infection were quantified using plaque assays on BHK-21 cells. Unpaired two-tailed t-test with Welch’s correction, p = 0.0002, t = 5.404, df = 11.81. (D) Specific Infectivity Ratios of progeny SINV were calculated as described earlier (1). Unpaired two-tailed t-test with Welch’s correction, p = 0.0084, t = 3.911, df = 5.820. For all panels error bars represent standard error of mean (SEM) of biological replicates and **P < 0.01; ****P < 0.0001.

We next assessed the effect of elevated *AMt2* expression on RNA virus infection by measuring the output of infectious progeny viruses following infection of cells expressing FLAG-*AMt2*. Ectopic MTase expression resulted in a four-fold increase in SINV titer, further supporting the positive *in vivo* correlation between *AMt2* expression and virus replication observed previously (Fig 2C). We also observed a concomitant increase in the per-particle infectivity of viruses upon assaying them on vertebrate baby hamster kidney fibroblast cells, as evidenced by higher specific infectivity ratios (Fig 2D). Together these results support the idea of *Aedes* DNMT2 being a proviral factor exploited by the virus to enhance its replication and transmission in the mosquito vector.

*AMt2* expression is reduced in the presence of *Wolbachia*. To test whether virus restriction *in vivo* is a consequence of reduced DNMT2 activity, we measured virus replication in mosquito cells following pharmacological inhibition of DNMT2. Structural homology of DNMT2 to other members of the DNA MTase family has allowed it to retain its DNA binding ability *in vitro*. However, they are canonically known to methylate tRNA molecules (24). Furthermore, DNMT2 is known to methylate RNA substrates by a different mechanism than canonical RNA methyltransferases. This mechanism of action makes DNMT2 susceptible to the action of DNA methyltransferase inhibitors ribo- (5-Azacytidine or 5-AZAC) or deoxyribo- (Deoxy-5-Azacytidine or DAC5) while ensuring that the function of other RNA methyltransferases in the cell remain unperturbed (25, 26). We reasoned that pretreatment of mosquito cells with either MTase inhibitor should reduce cellular DNMT2 activity and consequently restrict alphavirus replication. Pre-treating *Wolbachia-*free C710 cells with RNA MTase inhibitor 5-AZAC prior to infection reduced SINV RNA replication approximately 5-fold at 24 hours post-infection (Fig 3A). Virus titer was also reduced approximately 10-fold (Fig 3B). Finally, MTase inhibition also negatively influenced SINV per-particle infectivity, as evidenced by a 50-fold reduction in SI ratio (Fig 3C). Similar results were obtained for related alphavirus, Chikungunya virus (CHIKV, Fig S2A,B).

**Fig 3.**
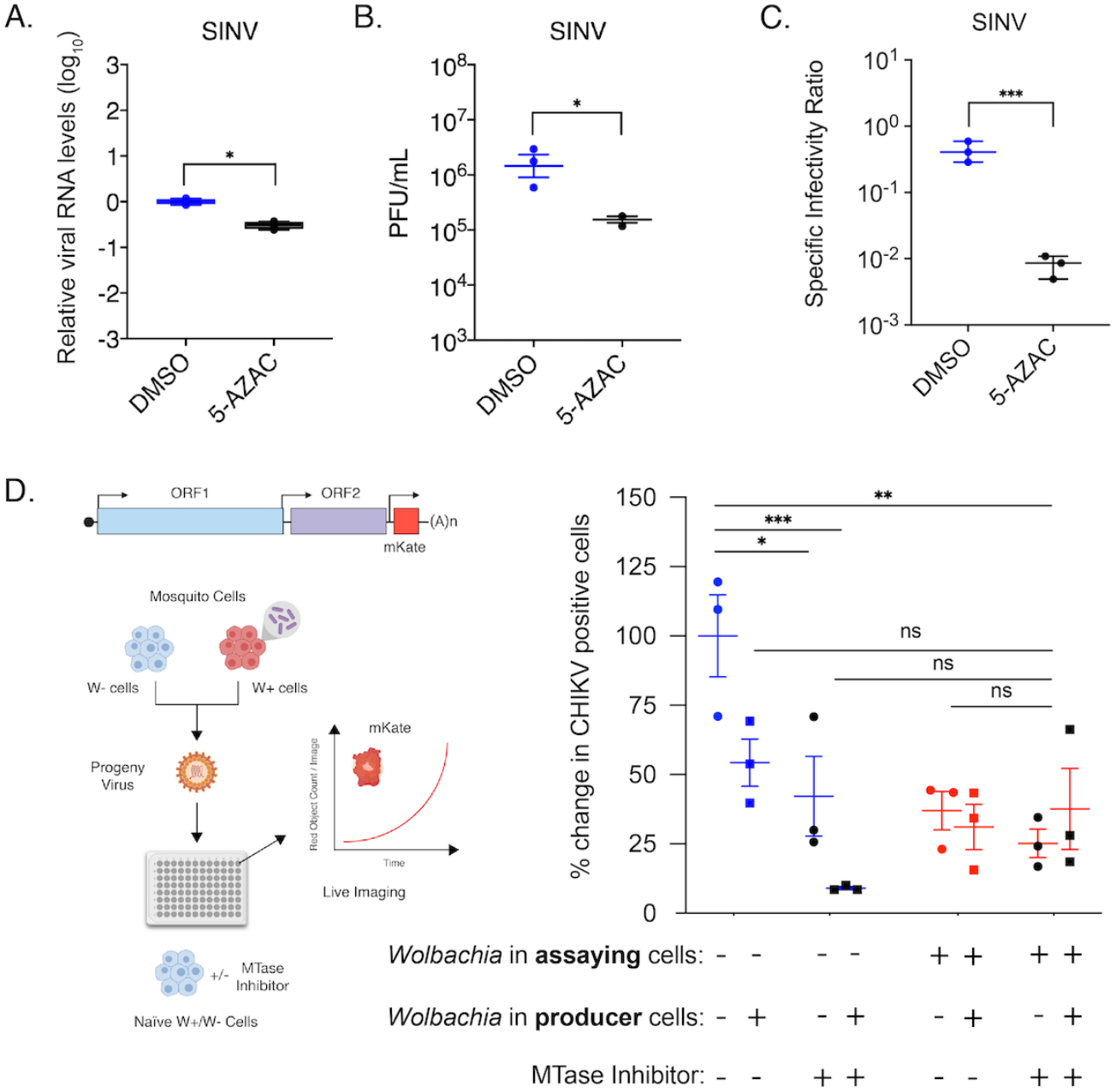
Pharmacological inhibition of mosquito DNMT2 reduces virus replication and per-particle infectivity. Inhibition of mosquito DNMT2 in *Wolbachia-free Aedes albopictus* derived C710 cells was carried out using MTase inhibitors 5-Azacytidine (5-AZAC). Dimethyl-sulfoxide (DMSO) was used as the negative control. In each case, cells were pretreated with 5 μM inhibitors overnight prior to infections with SINV at MOI of 10. Cell lysates and supernatants were harvested at 24 hours post infection to quantify cellular viral RNA levels and infectious titer, respectively. (A) Levels of SINV RNA in mosquito cells treated with MTase inhibitor 5-AZAC were determined using quantitative RT-PCR. Unpaired two-tailed t-test with Welch’s correction, SINV: p = 0.0012, t = 6.742, df = 4.892. (B) Infectious SINV titers produced from mosquito cells treated with MTase inhibitor 5-AZAC were determined using plaque assays on BHK-21 cells. Unpaired two-tailed t-test with Welch’s correction, SINV: p = 0.0339, t = 4.541, df = 2.322. (C) Specific infectivity ratios of progeny SINV was calculated as the ratio of infectious plaque forming units (B) over total viral genome copies present in collected cell supernatants as quantified by qRT-PCR. Unpaired two-tailed t-test with Welch’s correction, SINV: p = 0.0002, t = 12.59, df = 3.946. Error bars represent standard error of mean (SEM) of three independent experimental replicates. (D) CHIKV expressing mKate fluorescent reporter protein was grown in C710 *Aedes albopictus* cells in the presence (W+ virus) or absence (W− virus) of *Wolbachia* (strain *w*Stri). These progeny viruses were then used to infect C710 cells without and with *Wolbachia* (strain *w*Stri) pretreated with MTase inhibitor DAC5 (black data points) or DMSO (blue data points for *Wolbachia* free cells, red data points for *Wolbachia* colonized cells) synchronously at a MOI of 1 particle/cell. Virus growth in cells was measured in real time by imaging and quantifying the number of red cells expressing the virus encoded mKate protein forty-eight hours post infection, using live cell imaging. Shape of data points represent the origin of virus used to initiate infection; circles represent viruses derived from W− cells, boxes represent viruses derived from W+ cells. Three-way ANOVA with Tukey’s post hoc test for multivariate comparisons. For all panels error bars represent standard error of mean (SEM) of independent experimental replicates (n=3). *P < 0.05; **P < 0.01; ****P < 0.0001, ns = non-significant.

Using our previously published live-cell imaging system, we used a fluorescently-tagged CHIKV reporter virus (CHIKV-mKate) to examine the effect of deoxyribo- MTase inhibitor DAC5 on virus replication in *Wolbachia-free* and *Wolbachia-colonized Aedes albopictus* cells using the Incucyte live-cell imaging platform (1). As before, fluorescent protein expression was used as a proxy of virus replication in cells with (DAC5) and without (DMSO) inhibitor pretreatment. Virus replication was measured over 50 hours by quantifying mean virus-encoded red fluorescent reporter (mKate) expression observed over four distinct fields of view taken per well every 2-hours (Fig 3D, S2C).

In line with previous observations, viruses derived from *Wolbachia-*colonized cells (W+ virus, we will refer to viruses derived from *Wolbachia-*colonized cells as W+ and their counterparts, derived from *Wolbachia-*free cells as W− virus), produced under low *AMt2* conditions, are less infectious on W− cells, limiting their dissemination (1). Notably, this phenotype is particularly pronounced when W+ viruses encounter cells also colonized with the endosymbiont, presumably exhibiting reduced *AMt2* expression (Fig 3D, S2D-E). To further investigate the importance of DNMT2 activity in *Wolbachia-*mediated virus inhibition we treated cultured *Aedes albopictus* (C710) cells with DAC5, a DNMT2 inhibitor. We predicted that replication kinetics of W+ viruses in inhibitor-treated *Wolbachia-free* (W−) cells should phenocopy kinetics of W+ virus replication in *Wolbachia-*colonized (W+) cells. Additionally, the kinetics of W− virus replication in inhibitor-treated *Wolbachia-*free (W−) cells should phenocopy W− virus replication in *Wolbachia-*colonized (W+) cells (1). Three-way ANOVA was used to determine the effect of MTase inhibitor (DAC5), progeny virus type (derived from producer cells with or *Wolbachia-free*), and/or time on virus replication in recipient cells (assaying cells with or *Wolbachia-free*). In the presence of MTase inhibitor, replication of both W− and W+ viruses was reduced throughout infection, with a greater decrease in the replication of W+ viruses relative to W− viruses phenocopying the replication of W+ viruses in *Wolbachia-colonized* cells. Finally, replication of W− viruses in the presence of inhibitor was comparable to that of W+ viruses in mock-treated *Wolbachia-free* cells. We observed no synergistic effect of virus source and MTase inhibitor on virus replication in *Wolbachia-colonized* cells, likely due to low mean reporter activity. Altogether, these results demonstrate that alphavirus replication is negatively impacted by perturbed DNMT2 activity in either producer (W+ virus) or recipient mosquito cells (W− + DAC5 cells or W+ cells) and that the effect is compounded when both co-occur (W+ virus in W− + DAC5 cells or W+ virus in W+ cells).

### Ectopic DNMT2 expression rescues alphaviruses from *Wolbachia-*mediated inhibition

Evidence gathered indicates that *AMt2* downregulation is responsible for pathogen blocking in mosquitoes. Therefore, we ectopically overexpressed *AMt2* in *Wolbachia-colonized* mosquito cells to alleviate virus inhibition, including disruption of viral RNA synthesis and progeny virus infectivity (Fig 4A). We observed a significant reduction in viral RNA levels in *Wolbachia-colonized* cells relative to *Wolbachia-free* cells (Fig 4B). Interestingly, expression of *FLAG_Amt2* increased SINV RNA levels 70-fold in *Wolbachia-colonized* cells compared to cells carrying FLAG-empty vector, restoring virus RNA synthesis; One-way ANOVA Holm-Sidak’s multiple comparisons test, w/ Wolb vs. w/ Wolb + AMt2, p < 0.0001 (Fig 4B). Additionally, we observed a significant improvement in per-particle infectivity of progeny viruses derived from *Wolbachia-colonized* cells ectopically expressing *AMt2* (W+ AMt2+ virus, Fig 4C). Therefore, both phenotypes of pathogen blocking were abrogated upon *AMt2* over-expression. Given that endosymbiont titers can influence the degree of virus inhibition, we checked whether altering *AMt2* levels significantly impacted *Wolbachia* titer in cells. Quantitative PCR was used to measure relative *Wolbachia* titer in cells transfected with FLAG-AMt2 or FLAG-empty. However, no changes in endosymbiont titer were observed following ectopic *AMt2* expression (Fig 4D). Therefore, changes in *Wolbachia* titer do not explain the loss of pathogen blocking.

**Fig 4.**
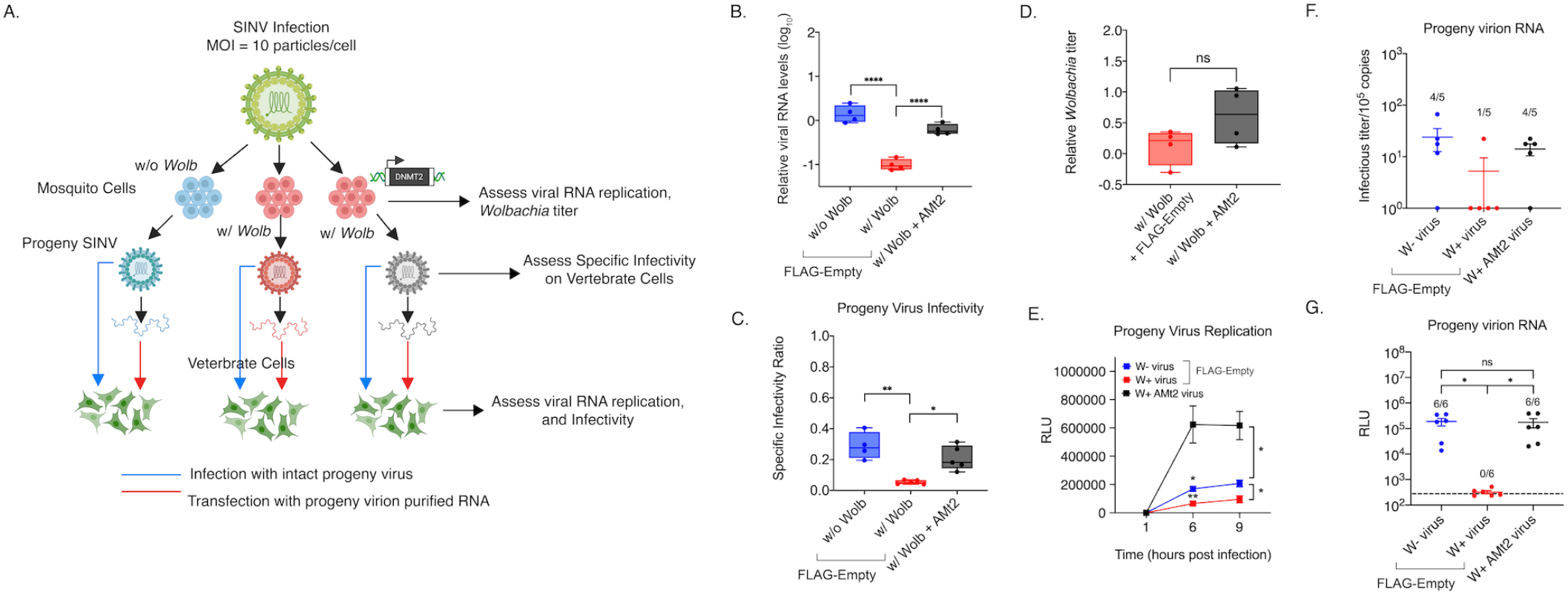
*AMt2* overexpression in *Wolbachia-*colonized cells rescues virus from endosymbiont-mediated inhibition. C710 cells with *Wolbachia* were transfected with expression vectors FLAG-empty (w/ Wolb) or FLAG-AMt2 (w/ Wolb + AMt2) for 48 hours prior to infection with SINV-nLuc at MOI of 10. *Wolbachia -*free cells expressing FLAG-empty (w/o Wolb) were used as a positive control. (A) Schematic of experimental workflow. (B) Viral genome replication in C710 cells was quantified using qRT-PCR using extracted total RNA from infected cell lysates. One-way ANOVA with Tukey’s post-hoc test of multivariate comparison. (C) Specific Infectivity Ratios of progeny viruses produced from the aforementioned infection was calculated as described earlier (1). Briefly, infectious progeny viruses collected from supernatants 48 hours post infection were quantified using plaque assays on BHK-21 cells, while total number of progeny virus particles was quantified via qRT-PCR of viral genome copies released into the supernatant. Error bars represent standard error of mean (SEM). One-way ANOVA with Tukey’s post-hoc test of multivariate comparison, w/ Wolb vs w/ Wolb + AMt2, p = 0.0003, w/o Wolb vs w/ Wolb, p < 0.0001. (D) C710 mosquito cells with *Wolbachia* were transfected with expression vectors FLAG-empty (w/ Wolb) or FLAG-AMt2 (w/ Wolb + AMt2) for 48 hours prior to quantification of endosymbiont titer via quantitative PCR using DNA from extracted cell lysates. Error bars represent standard error of mean (SEM). Unpaired, student’s t-test, p = 0.1316, t = 1.794, df = 5.097. Statistically non-significant values are indicated by ns. (E) Progeny viruses were used to synchronously infect naïve BHK-21 cells at equivalent MOIs of 5 particles/cell. Cell lysates were collected at indicated times post infection and luciferase activity (RLU), was used as a proxy for viral replication. Two-way ANOVA with Tukey’s post-hoc test of multivariate comparison, Time: p < 0.0001, *Wolbachia/AMt2*: p = 0.0003, Time x *Wolbachia/AMt2*: p < 0.0001. (F) Approximately 10^5^ copies (determined using qRT-PCR) each of virion encapsidated RNA extracted from the aforementioned W+, W+ AMt2 and W− viruses were transfected into naïve BHK-21 cells and infectious titer was determined by the counting the number of plaques produced after 72 hours post transfection. Numbers above bars refer to the proportion of samples that formed quantifiable plaque-forming units on BHK-21 cells. One-way ANOVA with Tukey’s post-hoc test of multivariate comparison. (G) 10^5^ copies each of virion encapsidated RNA extracted from the W+, W+ AMt2 and W− viruses were transfected into naïve BHK-21 cells and luciferase activity (RLU) was used as a proxy for viral replication at 9 hours post-transfection. Numbers above bars refer to the proportion of samples that produced luciferase signal above background levels, indicated by the dotted line. One-way ANOVA with Tukey’s post-hoc test of multivariate comparison, w/ Wolb vs w/ Wolb + AMt2: p < 0.00001, w/o Wolb vs w/ Wolb: p < 0.0001; w/o Wolb vs w/ Wolb + AMt2: p = 0.991. For all panels error bars represent standard error of mean (SEM). *P < 0.05; **P < 0.01; ****P < 0.0001.

Reduced per-particle infectivity of viruses occurring in the presence of *Wolbachia* (W+ viruses) is associated with reduced replication kinetics of these viruses in vertebrate cells and reduced infectivity of the encapsidated W+ virion RNA (1). As *AMt2* overexpression in *Wolbachia-colonized* mosquito cells rescued viral RNA synthesis and progeny virus infectivity, we examined the ability of progeny viruses derived from *Wolbachia-colonized* cells overexpressing *AMt2* (W+ derived AMt2+ cells) to replicate in vertebrate cells. We used SINV encoding a luciferase reporter for these experiments, allowing viral replication kinetics to determine the following synchronous infection of three progeny virus types: W− derived virus, W+ derived virus, and W+ derived AMt2+ virus. Replication of W+ derived AMt2+ viruses was significantly higher on a per-particle basis relative to W+ derived and, interestingly, W− derived viruses. This could be due to higher ectopic *AMt2* expression relative to what is induced natively during virus infection, implying perhaps a dose-dependent effect (Fig 4E). We then examined whether ectopic *AMt2* expression caused changes in the infectivity of the encapsidated virion RNA itself. Based on results from Fig 3D, we hypothesized that ectopic *AMt2* expression in *Wolbachia-colonized* cells should restore virion RNA infectivity. Indeed, following transfection of virion RNA into vertebrate BHK-21 cells, W+ derived virus RNA was largely non-infectious in contrast to RNA derived from viruses derived from W− and W+ AMt2+ cells (Fig 4F). Restored infectivity of W+ AMt2+ derived viral RNA was also validated using the luciferase-based virus replication assay (Fig 4G).

As demonstrated in Fig 2B, DNMT2 possesses the ability to bind viral RNA in mosquito cells. However, this alone does not indicate whether its MTase activity is essential for its proviral role. Broadly, DNMT2 comprises a catalytic domain and a target recognition domain responsible for RNA binding (27, 28). It is, therefore, possible that DNMT2’s regulatory role is independent of its MTase activity. To determine the importance of catalytic activity, we overexpressed a catalytically-inactive mutant of *AMt2*, replacing the highly conserved cysteine residue (C78) present in the motif IV region with a glycine (*AMt2* C78G, Fig 5A) in *Wolbachia-colonized* mosquito cells and asked whether this allele is capable of relieving pathogen blocking. Our data show *AMt2-*mediated rescue of SINV RNA synthesis and infectivity depends on its MTase activity as expression of the C78G mutant failed to rescue virus from *Wolbachia-mediated* inhibition (Fig 5B). We observed no improvement in SINV infectivity under these conditions (Fig 5C). Similar results were obtained from experiments carried out using CHIKV, where expression of wild-type *AMt2*, but not *AMt2-C78G*, resulted in improved virus titer (Fig 5D). However, while expression of *AMt2* increased CHIKV specific infectivity compared to expression of *AMt2-C78G* this increase was not statistically significant (Fig 5E). Based on these results, we conclude that the MTase *AMt2* promotes virus infection in mosquitoes and that lower *AMt2* expression in the presence of *Wolbachia* contributes to virus restriction and that MTase activity of DNMT2 is required for proviral function.

**Fig 5:**
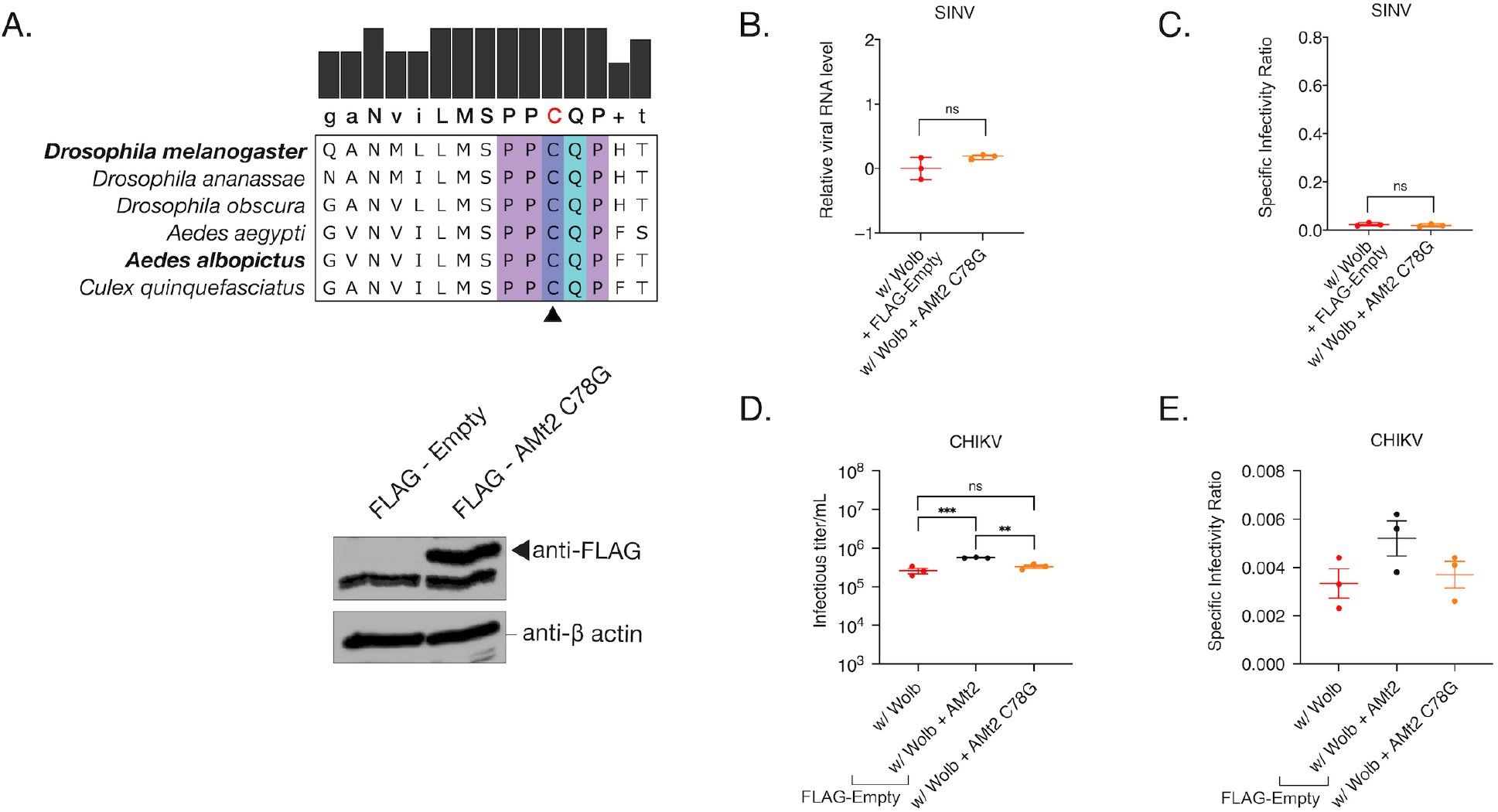
Catalytically inactive DNMT2 is unable to rescue *Wolbachia-mediated* virus inhibition in mosquito cells. (A) Multiple sequence alignment of the Motif IV region of DNMT2 derived from dipterans that are known to be colonized with native or non-native *Wolbachia*. The conserved catalytic cysteine (C) residue, depicted in red on the consensus sequence at the top, was mutated to a glycine (G) to abolish MTase activity of mosquito AMt2. Expression of the catalytic mutant (AMt2 C78G) was determined at 48 hours post transfection using Western Blot. C710 mosquito cells transfected with expression vector constructs with (FLAG-AMt2 C78G) or without (FLAG-empty) *AMt2*. Cytoplasmic lysates of cells were collected 48 hours post transfection and probed with anti-FLAG antibody. The non-specific band appearing below the desired size was used as a loading control. (B and C) C710 mosquito cells with *Wolbachia* were transfected with expression vectors FLAG-empty (w/ Wolb) or FLAG-C78G AMt2 (w/ Wolb + AMt2 C78G) for 48 hours prior to infection with SINV at MOI of 10. Infectious virus and Specific Infectivity (SI) of progeny viruses produced after 72 hours post infection were quantified as before. Unpaired t-test with Welch’s correction, p = 0.1734, t = 1.920, df = 2.396 (B), p = 0.4544, t = 0.8291, df = 3.937 (C). (D and E) C710 mosquito cells with *Wolbachia* were transfected with expression vectors FLAG-empty (w/ Wolb), FLAG-AMt2 (w/ Wolb + AMt2) or FLAG-C78G AMt2 (w/ Wolb + AMt2 C78G) for 48 hours prior to infection with CHIKV-mKate at MOI of 10. Infectious virus (D) and Specific Infectivity (E) of progeny viruses produced after 72 hours post infection were quantified as before. One-way ANOVA followed by Tukey’s post hoc test for multivariate comparisons for (D), w/ Wolb vs w/ Wolb + AMt2, p = 0.0009, w/ Wolb vs w/ Wolb + AMt2 C78G p = 0.2694, w/ Wolb + AMt2 vs w/ Wolb + AMt2 C78G p < 0.0040. Error bars represent standard error of mean (SEM) of biological replicates. **P < 0.01; ***P < 0.001, ns = non-significant.

### DNMT2 orthologs from mosquitoes and fruit flies regulate virus infection differentially in their respective hosts

The proviral role of the *AMt2* is intriguing, given the previously described antiviral role for the corresponding fruit fly ortholog, *Mt2* (9, 29). Interestingly, in our previous study, we observed that knocking down *Mt2* led to increased progeny virus infectivity. Therefore, we reasoned that ectopic expression of *Mt2* should reduce Sindbis virus infectivity (Fig S3). As with mosquito *AMt2*, we asked whether this involved direct targeting of viral RNA. AZA-IP of epitope-tagged Mt2 confirmed direct interactions between viral RNA and fly DNMT2 in *Wolbachia-free Drosophila melanogaster-*derived JW18 cells, which showed a 10-fold enrichment in SINV RNA-binding relative to a control host transcript (18S) (Fig S3A, B).

In contrast to the proviral effect of mosquito *AMt2*, ectopic *Mt2* expression significantly reduced infectivity of progeny SINV and CHIKV (W− derived Mt2+ virus) relative to those produced from cells expressing the control vector (W− derived virus), confirming our previous findings (Fig S3C). As with *AMt2*, we assessed whether reduced infectivity of W− derived Mt2+ viruses was due to their inability to replicate in vertebrate cells. Indeed, results from our luciferase reporter based viral replication assay revealed significantly reduced replication of W− Mt2+ derived viruses relative to W− derived viruses in vertebrate BHK-21 cells, similar to the behavior observed for W+ derived viruses (Fig S3D). Finally, we quantified the infectivity of virion encapsidated RNA from W− derived Mt2+ SINV and CHIKV viruses by measuring the number of plaque-forming units generated following transfection into vertebrate BHK-21 cells. For both SINV and CHIKV, the infectivity of virion encapsidated RNA was reduced for W+ viruses. Notably, this was phenocopied by virion RNA isolated from W− Mt2+ derived SINV and CHIKV (Fig S3E,F).

Similar to mosquito *AMt2*, fly *Mt2’s* ability to regulate virus fitness also rely on its catalytic activity, as expressing a catalytically inactive mutant (*Mt2* C78A) was unable to restrict the production of infectious virus and per-particle infectivity of SINV (Fig S4). Taken together, these results suggest that progeny virus/virion RNA infectivity is reduced in fly cells under conditions where MTase expression is elevated natively in the presence of *Wolbachia* (W+ virus) or artificially (W− Mt2+ virus).

### The presence of *Wolbachia* in mosquito cells is associated with altered viral RNA methylation

That ectopic *AMt2* expression in *Wolbachia-colonized Aedes albopictus* cells can restore the infectivity of SINV progeny virion RNA implies two important things; (i) Sindbis virion RNA carries 5-methylcytosine (m5C) modifications, and (ii) that altered *AMt2* expression in the presence of *Wolbachia* is associated with changes in the overall m5C content of the virion RNA. To directly determine if virus RNA is modified differentially in the presence of *Wolbachia*, we subjected virion RNA isolated from progeny SINV produced from *Aedes albopictus* cells colonized with (W+ virus) and without (W− virus) *Wolbachia* to liquid chromatography tandem mass spectrometry (LC-MS/MS) analyses (Fig 6A). We chose to focus our efforts on identifying the presence of 5-methylcytosine (m5C) and 6-methyladenosine (m6A) residues on the viral genome for our present analyses. We examined m6A due to recent reports highlighting the importance of this modification in regulating RNA virus replication (30, 31). A potential complication for these analyses is the presence of residue(s) of similar mass to charge ratio(s) to m5C, such as m3C. However, as shown in Fig S5A, we observed distinct distribution of the individual m3C and m5C peaks in the spectral output, demonstrating our ability to distinguish between these two bases (Fig S5A).

**Figure 6:**
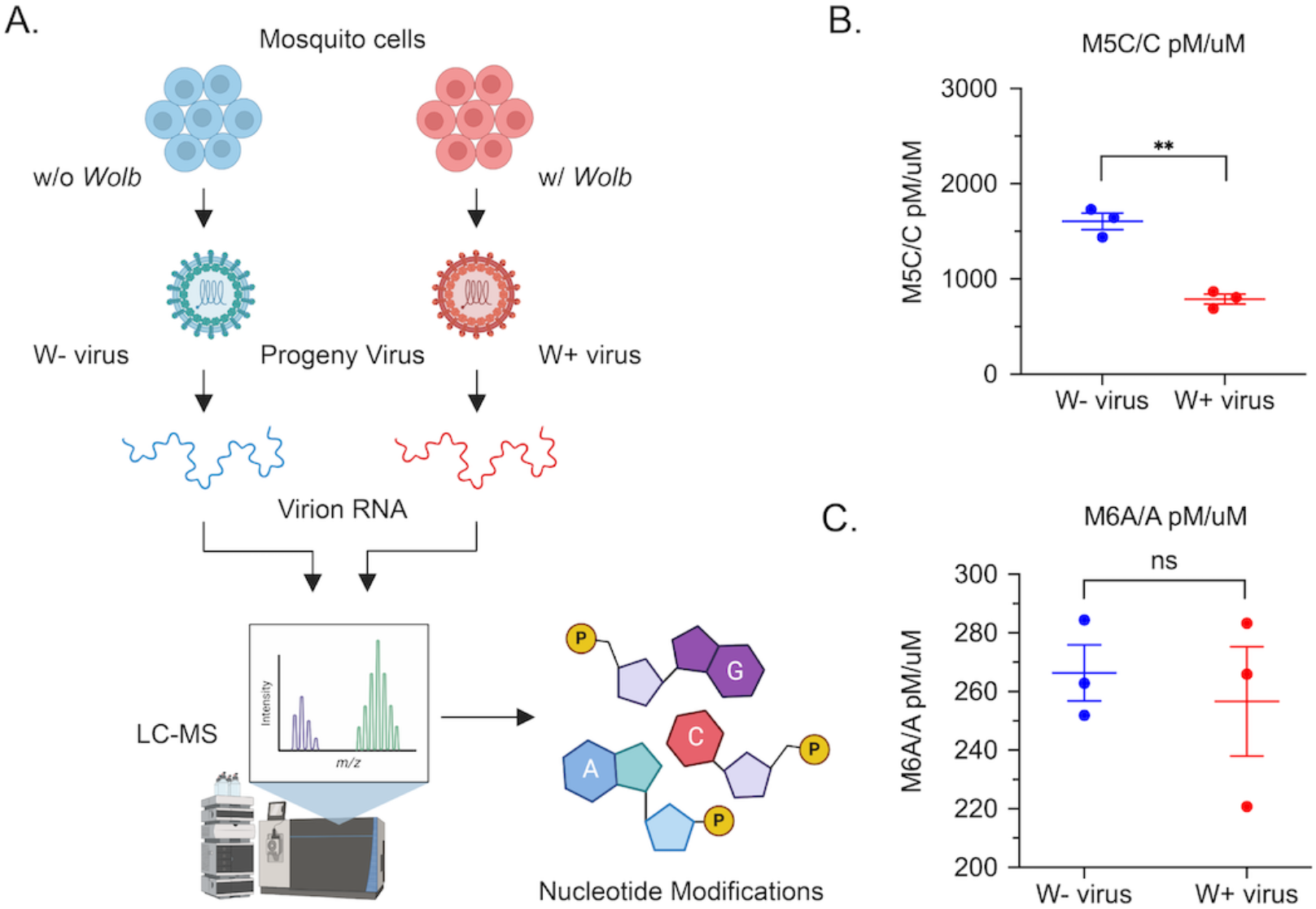
Presence of *Wolbachia* is associated with altered virion RNA methylation. (A) RNA isolated from progeny viruses derived from mosquito cells colonized with (W+ virus) or without (W− virus) *Wolbachia* were subjected to LC-MS/MS analyses to determine their nucleotide content. (B) Normalized 5-methyl cytosine (M5C) content of RNA isolated from W− and W+ viruses represented as a ratio of total unmodified cytosine content. Unpaired two-tailed t-test, p = 0.0013, t = 8.080, df = 4 (C) Normalized 6-methyl adenosine (M6A) content of RNA isolated from W− and W+ viruses represented as a ratio of total unmodified adenosine content. Unpaired two-tailed t-test, p = 0.666, t = 0.4643, df = 4. Error bars represent standard error of mean (SEM) of three independent virus preps from each cell type. **P < 0.01; ns = non-significant.

LC-MS/MS analyses of RNA purified from virion RNA derived from *Wolbachia* free (W−) and *Wolbachia*-colonized (W+) cells demonstrated W+ virion RNA to contain, on average, more than 2-fold fewer m5C residues compared to W− virion RNA across three independent virus preps from each cell type (Fig 6B). Notably, both W+ and W− virion RNA was determined to consist of comparable levels of m6A residues across all biological replicates (Figure 6C). In addition, we observed no significant changes in the overall m3C content between W+ and W− virion RNA (Fig S5B). It should be noted that while we did not observe changes in the overall abundance of m6A and m3C residues between W+ and W− virion RNA, it is unclear whether the presence of *Wolbachia* leads to altered distribution of m6A and/or m3C modifications in the context of the overall SINV RNA sequence. Finally, we used LC-MS/MS analyses to quantify viral Type-0 (7-methyl-GpppNp or M7G) cap structures present in W− and W+ virion RNA to estimate relative ratios of capped versus non-capped virus progeny produced in the presence or absence of *Wolbachia*. While there were no statistically significant differences present between the respective W− and W+ sample means (t-test: p=0.999), we found M7G content to vary significantly among W+ virion RNA replicates (F-test: p=0.0088), indicating either that the ratios of capped vs. non-capped viruses vary significantly within virus populations derived from *Wolbachia-colonized* cells, or that viral RNAs produced under these conditions carry varying amounts of internal M7G signatures (Fig S5C).

These data are consistent with: (i) DNMT2 being an essential host factor in mosquito cells for efficient virus replication and transmission, (ii) this proviral effect being exert through m5C modification of the viral genomic RNA, and (iii) a mechanism of *Wolbachia-*mediated pathogen blocking being the reduction of DNMT2 expression.

## Discussion

Virus inhibition in *Wolbachia*-colonized arthropods is associated with two distinct features independent of any particular host-*Wolbachia* strain combination; (i) reduced genome replication of the +ssRNA viruses in *Wolbachia*-colonized cells, and (ii) reduced per-particle infectivity of progeny +ssRNA viruses produced under these conditions (1). While these shared attributes constitute a subset of several virus inhibition phenotypes, it indicates the existence of a conserved cellular mechanism of restriction. In our previous study, we used the prototype alphavirus, Sindbis as our +ssRNA virus model to uncover an essential role of the fruit fly RNA cytosine methyltransferase (MTase) gene *Mt2* (DNMT2) as an essential host determinant of *Wolbachia-*mediated pathogen blocking (9). Furthermore, loss of *Drosophila* DNMT2 is associated with a loss in virus inhibition by *Wolbachia* and increased progeny virus infectivity in mammalian cells, suggesting that DNMT2 might regulate these two aspects of alphavirus replication. These findings thus led us to ask the following question in our present study: Is DNMT2 a conserved host determinant of *Wolbachia-mediated* +ssRNA virus inhibition between fruit flies and mosquitoes?

MTase expression in adult *Aedes aegypti* mosquitoes is distinctly altered in the presence of both virus and *Wolbachia* and in opposite directions (Fig 1). DNMT2 expression is elevated in the body of mosquitoes following an infectious bloodmeal. Notably, this pattern is observed in the salivary gland tissues, representing the final site of virus production in the vector prior to transmission to a vertebrate. We show that this is beneficial to the virus, as ectopic MTase expression in cultured, *Wolbachia-*free *Aedes albopictus* mosquito cells promotes virus replication and, importantly, progeny virus infectivity. This has also been reported for DENV-2 infection in *w*Mel-colonized *Aedes aegypti* mosquitoes (22). We observed that baseline MTase activity is required for virus replication and spread in *Aedes* cells (Fig S2). Furthermore, the extent to which virus replication is affected by MTase inhibitors depends on the virus source, with viruses produced from *Wolbachia-*colonized cells (W+ viruses) being more susceptible to MTase inhibition.

This outcome phenocopies the scenario in which virus spread is most restricted under conditions where both producer and target mosquito cells are colonized with *Wolbachia* (1). In line with these findings, our data indicate a decrease in MTase expression occurs in the presence of *Wolbachia* (Fig 1). This observation is in line with previous reports (22). Thus, our collective data support a model in which endosymbiont-dependent inhibition of MTase expression and catalytic MTase function contribute to reduced virus replication and per-particle infectivity in mosquitos (Fig 2-4). This consequently limits virus dissemination within the vector and transmission to a vertebrate host (Fig 2-4). Given that our results are coherent with prior reports involving a different RNA virus and *Wolbachia* strain, the interaction between virus, *Wolbachia*, and host MTase expression is likely independent of any particular virus-host-*Wolbachia* combination, representing a conserved feature of pathogen blocking in the native *Aedes* vector.

Our data demonstrate an interaction between DNMT2 orthologs from *Aedes albopictus* and *Drosophila melanogaster* and viral RNA (Fig 5B, Fig S4B) (1). However, it remains to be seen whether these interactions are analogous to DNMT2-DCV RNA interactions in *Drosophila*, where the MTase-viral RNA binding occurs specifically at structured viral Internal Ribosomal Entry Sites (IRES) (29). Additionally, it is unclear if DNMT2 interactions in the mosquito cell are specific for viral RNA or whether it extends to host transcripts. Future studies involving PAR-CLIP-sequencing of immunoprecipitated DNMT2-RNA complexes should allow the identification and mapping of distinct DNMT2-binding motifs and/or structural elements within viral and host RNA species. Nevertheless, sites of DNMT2 recruitment to viral RNA, and viral and host proteins are required for recruitment remain unidentified. Assuming that the proviral role of *Aedes* DNMT2 involves the addition of m5C signatures to specific residues on the viral genome, it seems likely that viral co-factor(s) are required for specificity. *Drosophila* DNMT2 is antiviral in fruit flies, however *Drosophila* is not a natural host for alphaviruses, and it is likely that the adaptation of the virus to the natural vector has facilitated an appropriate proviral interaction with *Aedes* DNMT2 or an associated *Aedes-*specific host factors(s) absent in *Drosophila melanogaster* (32).

The m5C content of virion RNA produced from *Wolbachia-colonized* cells (W+ viruses) is significantly reduced relative to cells without the symbiont (Fig 6B) (1), consistent with DNMT2s role as a cytosine MTase. Incidentally, this finding follows reports dating back several decades describing the occurrence of m5C residues within intracellular SINV RNA (33). Relative abundance of methylated to total cytosine residues on SINV virion RNA derived from *Wolbachia-*free mosquito cells is 15:10000, accounting for cytosine content in the viral genome we predict there to be 4 to 5 m5C signatures per encapsidated virion RNA genome produced in *Aedes albopictus* cells. Our observation supports the involvement of these intracellular m5C signatures in alphavirus genome replication that W+ virion RNA, which are presumably hypomethylated, are less infectious on a per-genome basis (Fig 4E-F) (34). Indeed, based on our data, we can infer that these m5C modifications regulate alphavirus infection across multiple hosts and, thus, by extension, aspects of the virus transmission cycle. It should also be noted that while methylated nucleotide residues like m6A and m5C occur on RNA virus genomes at higher rates than those present in cellular RNA species, our results do not exclude the possibility of other RNA modifications, as well as differential modification of host RNA species and playing a role in regulating virus replication and transmission. This may be of particular consequence given recent evidence of altered m6A modification of specific cellular transcripts during flavivirus infection in vertebrate cells (31, 35, 36).

Alphaviruses derived from mosquito cells are more infectious on vertebrate cells on a per-particle basis than vertebrate cell-derived viruses and vice versa (37). This carries the implication that progeny viruses originating from one cell type may possess intrinsic properties that can confer a fitness advantage while infecting a destination host cell type, altering their infectivity on these destination cells on a per-particle level. As to what such properties may represent, current evidence points towards differences in virus structure, such as differential sialation or glycosylation of viral glycoproteins impacting host receptor-binding and/or differences in the encapsidated cargo, e.g., packaging of host ribosomal components (38–40). Furthermore, as our results suggest, another property that might confer unique cell-type-specific advantages to viruses is the differential modification of the virion RNA. Indeed, recent evidence shows that modifications like N^6^-methyladenosine (m6A) and 5-methylcytosine (m5C) can regulate viral RNA functionality in the cell (30, 41). Therefore, it is possible that such modifications also influence the infectivity of progeny viruses produced from said cells. However, how these modifications affect virus replication in a cell is still an open question. Indeed, information regarding the functional consequence of m5C or other RNA modifications on viral RNA is limited. Thus, while we may draw certain conclusions based on our current knowledge of known eukaryotic RNA modifications, the potential implications of arbovirus RNA methylation may be broader than we are currently able to anticipate (42). We can hypothesize that differential viral methylation may alter host responses to infection, in that depending on the host or cell type, as well as the genomic context of methylation, presence or absence of m5C may either allow detection by and/or provide a mechanism of escape from RNA-binding proteins (e.g., Dicer, RIG-I, MDA5, TLRs, APOBEC3) involved in virus restriction or non-self RNA recognition that trigger downstream immune signaling and interferon production (43). Differential modifications of viral RNA may thus also regulate different cytological outcomes of arboviruses infection of arthropod and vertebrate cells i.e. persistence versus cell death.

It remains to be seen whether or not one or more of these situations occur during pathogen blocking and if W− and W+ viruses trigger differential innate immune responses in vertebrate cells. Based on our data, we propose a model in which our current estimates of m5C residues on W− viruses represent the “wild-type” epitranscriptome of mosquito-derived alphavirus. In naïve vertebrate cells, the presence of these signatures allows viruses to replicate efficiently following successful evasion of host innate immunity. In contrast, m5C hypomethylation of W+ viruses renders them more susceptible to host-induced restriction, thus impacting their ability to propagate. Aside from heightened immune susceptibility, the decreased fitness of hypomethylated W+ viruses could also result from reduced incoming viral RNA stability and/or translation.

Given that pharmacological inhibition of MTase activity impacts virus spread in mosquito cells, it is likely that W+ virus hypomethylation also influences dissemination in arthropod cells (1). However, it is also possible that other factors contribute to the reduced fitness of W+ viruses. In particular, our LC-MS/MS analyses suggest increased heterogeneity in m7G moiety abundance on W+ virion RNA, indicating either difference in abundance of internal m7G methylation signatures or a potential imbalance in viral RNA capping in the presence of *Wolbachia*. In addition, past work has shown that SINV populations derived from different hosts vary with regard to the ratios of capped and non-capped SINV RNA (44). Despite being important for alphavirus replication, non-capped SINV RNA alone are compromised in their ability to undergo translation, are more susceptible to RNA decay machineries, and have been shown to induce elevated innate immune response, all of which might contribute to the observed loss in infectivity.

Finally, the data presented here implicating epitranscriptomic regulation of alphaviruses unlocks multiple avenues of investigation, which include, but are not limited to the following. First, it is crucial to determine the genomic context of m5C and other RNA modifications on viral RNA for different hosts, cell types, and infection timeline, which may be achieved by long-read, direct RNA sequencing from virus-infected cells. Doing so would allow sequence-specific mapping of these signatures and help address whether virus infection is regulated solely via targeting viral RNA by cellular MTases. Furthermore, deriving mapping information might inform us whether modifications are directed to specific RNA elements that result in spatiotemporal changes in RNA structure and altered base-pairing, thus regulating virus RNA polymerase fidelity and/or translation in the cell. Additional areas of inquiry involve identifying cellular pathways responsible for determining the fate of W+ viruses and characterizing the functional consequences of abolishing highly conserved m5C residues on the viral RNA. This would allow further exploration into the effect of these signatures on RNA stability, gene expression, and/or packaging across arthropod and vertebrate cells. Lastly, unlike m6A-modifications, little is known regarding how m5C signatures are interpreted, i.e., how they are “read,” “maintained,” and “erased,” in mammalian and, to an even lesser extent, in arthropod cells (42). Promising candidates include m5C-binding “reader” proteins ALYREF and YBX1, which function alongside the known cellular m5C MTase NSUN2 to influence mRNA nuclear transport and stability (45, 46). Following approaches described in recent studies, identification of these RNA-binding proteins, either viral or host-derived, may be achieved via affinity-based immunoprecipitation of viral RNA and form the basis of future studies (47).

Like most other RNA viruses, Alphaviruses are limited in their coding capacity and are known to alter their genome structure under various cellular conditions to regulate aspects of their replication as a way to maximize viral genome functionality. Echoing this idea, the findings presented in this study add to our understanding of regulatory mechanisms adopted by these viruses to successfully navigate within and transition between vertebrate and arthropod host species.

## Materials and Methods

### Insect and Mammalian Cell Culture

RML12 *Aedes albopictus* cells with and *Wolbachia-free w*Mel was grown at 24 °C in Schneider’s insect media (Sigma-Aldrich) supplemented with 10% heat-inactivated fetal bovine serum (Corning), 1% each of L-Glutamine (Corning), non-essential amino acids (Corning), and penicillinstreptomycin-antimycotic (Corning). C710 *Aedes albopictus* cells with and *Wolbachia-free* were grown at 27 °C under 5% ambient CO_2_ in 1X Minimal Essential Medium (Corning) supplemented with 5% heat-inactivated fetal bovine serum (Corning), 1% each of L-Glutamine (Corning), non-essential amino acids (Corning) and penicillin-streptomycin-antimycotic (Corning). Vertebrate baby hamster kidney fibroblast or BHK-21 cells were grown at 37 °C under 5% ambient CO2 in 1X Minimal Essential Medium (Corning) supplemented with 10% heat-inactivated fetal bovine serum (Corning), 1% each of L-Glutamine (Corning), non-essential amino acids (Corning) and penicillin-streptomycin-antimycotic (Corning). JW18 *Drosophila melanogaster* cells with and without *Wolbachia w*Mel were grown at 24 °C in Shields, and Sang M3 insect media (Sigma-Aldrich) supplemented with 10% heat-inactivated fetal bovine serum, 1% each of L-Glutamine (Corning), non-essential amino acids (Corning), and penicillin-streptomycin-antimycotic (Corning).

### Mosquito rearing and blood meals

*Aedes aegypti* mosquitoes either -infected and -uninfected with *Wolbachia* (*w*AlbB strain) (generously provided by Dr. Zhiyong Xi, Michigan State University, USA), were reared in an insect incubator (Percival Model I-36VL, Perry, IA, USA) at 28 °C and 75% humidity with 12 h light/dark cycle. Four to six-day-old, mated female mosquitoes were allowed to feed for one hour on approximately 10^8^ PFUs of SINV (TE12-untagged) containing citrated rabbit blood (Fisher Scientific DRB030) supplemented with 1mM ATP (VWR), and 10% sucrose using a Hemotek artificial blood-feeding system (Hemotek, UK) maintained under a constant temperature of 37 °C. Engorged mosquitoes were then isolated and reared at 28 °C in the presence of male mosquitoes. For harvesting whole tissues, mosquitoes were harvested 5-7 days post blood meal before being snap-frozen in liquid nitrogen and stored at −80 °C before further processing. For salivary gland dissections, mosquitoes were kept immobilized on ice before dissection. Collected salivary gland tissues were washed three times in a cold, sterile saline solution (1XPBS) before being snap-frozen in liquid nitrogen and stored at −80 °C before further processing. Three salivary glands were pooled to create each biological replicate. Samples for qPCR and qRT-PCR were homogenized in TRiZOL (Sigma Aldrich) reagent and further processed for nucleic acid extractions using manufacturer’s protocols.

### Virion RNA extraction and transfection

Virion encapsidated RNA was extracted from viruses (SINV-nLuc) were purified over a 27% sucrose cushion using TRiZOL reagent (Sigma Aldrich) using the manufacturer’s protocol. Post extraction, RNAs were DNase (RQ1 RNase-free DNase, NEB) treated according to the manufacturer’s protocol to remove cellular contaminants, and viral RNA copies were quantified via quantitative RT-PCR using primers probing for SINV nsP1 and E1 genomic regions (Table S1) and a standard curve comprised of linearized SINV infectious clone containing the full-length viral genome. To determine infectivity or replication kinetics of Sindbis virion RNA, equal copies of virion isolated RNA (10^5^ copies), quantified using qRT-PCR, were transfected into BHK-21 cells in serum-free Opti-MEM (Gibco). Transfection was carried out for 6 hours before the transfection inoculum was removed, and overlay was applied. Cells were fixed post-transfection using 10% (v/v) formaldehyde and stained with crystal violet to visualize plaques. To maximize the production of infectious units, equal mass (1 μg) of virion (SINV-nLuc) isolated RNA derived from JW18 fly cells was transfected into BHK-21 cells. Transfection was carried out for 6 hours before the transfection inoculum was removed, and overlay was applied. Cells were fixed 48 (SINV) or 72 (CHIKV) hours post-transfection using 10% (v/v) formaldehyde and stained with crystal violet to visualize plaque-forming units.

### Viral replication assays

The viral genome and sub-genome translation were quantified using cellular lysates following synchronized infections with reporter viruses (SINV-nLuc) or transfections with virion-derived RNA from the aforementioned viruses. At indicated times post-infection, samples were collected and homogenized in 1X Cell Culture Lysis Reagent (Promega). In addition, samples were mixed with NanoGlo luciferase reagent (Promega), incubated at room temperature for three minutes before luminescence was recorded using a Synergy H1 microplate reader (BioTech instruments).

### Virus infection in cells and progeny virus production

Virus stocks were generated from RML12, C710, or JW18 cells, either with or without *Wolbachia* or overexpressing DNMT2 orthologs by infecting naïve cells with the virus at an MOI of 10. In all cases, serum-free media was used for downstream virus purification. Media containing virus was collected five days post-infection for alphaviruses SINV (SINV-nLuc, TE12-untagged, TE3’2J-GFP, and TE3’2J-mCherry) and CHIKV (CHIKV18125-capsid-mKate). Virus stocks were subsequently purified and concentrated by ultracentrifugation (43K for 2.5 h) over a 27% (w/v) sucrose cushion dissolved in HNE buffer. Viral pellets were stored and aliquoted in HNE buffer before being used for all subsequent experiments.

### DNMT2 overexpression in arthropod cells

*Aedes albopictus AMt2* coding region was subcloned into PCR 2.1 TOPO vector (Invitrogen) by PCR amplification of cDNA generated using reverse transcribed from total cellular RNA isolated from C636 *Aedes albopictus* cells using Protoscript II RT (NEB) and oligo-dT primers (IDT). The coding region was validated via sequencing before being cloned into the pAFW expression vector (1111) (Gateway Vector Resources, DGRC), downstream of and in-frame with the 3X FLAG tag using the native restriction sites *AgeI* and *NheI* (NEB). Expression of both FLAG-tagged AaDNMT2 in mosquito cells was confirmed using qRT-PCR and Western Blots using an anti-FLAG monoclonal antibody (SAB4301135 - Sigma-Aldrich, 1:3000 dilution in 2% Milk in 1X TBS + 1% Tween-20) (Fig 4A). In addition, catalytic MTase mutant of *AMt2* (*AMt2-C78G*) was generated via site-directed mutagenesis (NEB, Q5 Site-Directed Mutagenesis Kit), using primers listed in the primer table (Table S1). *Drosophila Mt2* (FBgn0028707) cDNA clone (GM14972) obtained from DGRC (https://dgrc.bio.indiana.edu/) was cloned into the pAFW expression vector (1111) with an engineered *SaII* site (Gateway Vector Resources, DGRC) downstream of and in-frame with the 3X FLAG tag using Gibson assembly (HiFi DNA assembly mix, NEB). Expression of FLAG-tagged DNMT2 in fly cells was confirmed using qRT-PCR and Western Blots using an anti-FLAG monoclonal antibody (SAB4301135 - Sigma-Aldrich, 1:3000 dilution in 2% Milk in 1X TBS + 1% Tween-20). Catalytically inactive *Mt2* (*Mt2* C78A) variant was generated via site-directed mutagenesis (NEB, Q5 Site-Directed Mutagenesis Kit) using primers listed in the primer table (Table S1).

### Immunoprecipitation of DNMT2-viral RNA complexes

JW18 fly cells and C710 mosquito cells were transfected with expression vectors FLAG-Mt2 and FLAG-AMt2 respectively for 48 hours before infection with SINV at MOI of 10. Control cells were transfected with the empty vector plasmid FLAG-empty. In addition, cells were treated for approximately 18h with 5 μM 5-Azacytidine to covalently trap Mt2 or AMt2 with its target cellular RNA before RNA immunoprecipitation using an anti-FLAG antibody following manufacturer’s protocols (SAB4301135 - Sigma-Aldrich, 1:100 dilution) (23).

### Real-time quantitative RT-PCR analyses

Following total RNA extraction using TRiZOL reagent, cDNA was synthesized using MMuLV Reverse Transcriptase (NEB) with random hexamer primers (Integrated DNA Technologies). Negative (no RT) controls were performed for each target. Quantitative RT-PCR analyses were performed using Brilliant III SYBR Green QPCR master mix (Bioline) with gene-specific primers according to the manufacturer’s protocol and the Applied Bioscience StepOnePlus qRT-PCR machine (Life Technologies). The expression levels were normalized to the endogenous 18S rRNA expression using the delta-delta comparative threshold method (ΔΔCT). Fold changes were determined using the comparative threshold cycle (CT) method (Table S1). Efficiencies for primer sets used in this study have been validated in our previous study (1).

### DNMT2 inhibition in mosquito cells

Inhibition of *Aedes* DNMT2 activity in C710 cells was achieved using RNA and DNA cytosine methyltransferase inhibitors, 5-aza-cytidine (5-AZAC, Sigma-Aldrich) and 5-deoxy-azacytidine (DAC-5, Sigma-Aldrich). In each case, *Aedes albopictus* C710 cells were treated overnight with media containing either 5μM inhibitor diluted in Dimethyl sulfoxide (DMSO) or DMSO alone. Due to the poor stability of 5-AZAC, media containing fresh inhibitors was added every day post-infection (48).

### Live cell imaging

Live-cell imaging experiments were carried out using a setup similar to our previous study (1). The growth of fluorescent reporter viruses in *Aedes albopictus* (C710) cells was monitored using Incucyte live-cell analysis system (Essen Biosciences, USA). *Aedes albopictus* C710 cells were grown under standard conditions as described earlier under 5% ambient CO_2_ at 27 °C. Cells were plated to 75-80% confluency in 96-well plates to separate adjacent cells and preserve cell shape for optimal automated cell counting. To synchronously infect cells virus was adsorbed at 4°C. Post adsorption, cell monolayers were extensively washed with cold 1XPBS to remove any unbound virus particles, followed by the addition of warm media (37°C) to initialize virus internalization and infection. Cells per well were imaged and averaged across four distinct fields of view, each placed in one-quarter of the well every 2 hours throughout the infection. Total fluorescence generated by cells expressing the red fluorescent reporter mKate was calculated and normalized by the cell number for every sample. A manual threshold was set to minimize background signal via automated background correction at the time of data collection. Following the acquisition, data were analyzed in real-time using the native Incucyte ^®^ Base Analysis Software.

### Quantification of RNA modification by LC-MS/MS

Total RNA (3-7μg) was digested by nuclease P1 (10 Units) at 50°C for 16 hr. Additional Tris pH 7.5 was then added to a final concentration of 100 mM to adjust, followed by the addition of calf intestinal alkaline phosphatase (CIP, NEB, 2Units). The mixture was incubated at 37°C for 1 hour to convert nucleotide 5’-monophosphates to their respective nucleosides. Next, 10μl of RNA samples were diluted to 30 μL and filtered (0.22 μm pore size). 10μL of the sample was used for LC-MS/MS. Briefly, nucleosides were separated on a C18 column (Zorbax Eclipse Plus C18 column, 2.1 x 50mm, 1.8 Micron) paired with an Agilent 6490 QQQ triple-quadrupole LC mass spectrometer using multiple-reaction monitoring in positive-ion mode. The nucleosides were quantified using the retention time of the pure standards and the nucleoside to base ion mass transitions of 268.1 to 136 (A), 244.1 to 112 (C), 284.2 to 152 (G), 258 to 126 (m3C and m5C), 282.1 to 150 (m6A), 298 to 166 (m7G). Standard calibration curves were generated for each nucleoside by fitting the signal intensities against concentrations of pure-nucleoside preparations. The curves were used to determine the concentration of the respective nucleoside in the sample. The A, G, and C standards were purchased from ACROS ORGANICS; m5C was purchased from BioVision; m7G, m1G, and m3C were purchased from Carbosynth, m6G and m6A were purchased from Berry’s Associates, and m1A was from Cayman Chemical Company. The modification level on the nucleosides was calculated as the ratio of modified: unmodified.

## Supporting information

all supplemental material

## Data availability

The full Incucyte dataset is available in the form of an excel sheet labeled Dataset S1.

## Statistical analyses of experimental data

All statistical analyses were conducted using GraphPad Prism 8 (GraphPad Software Inc., San Diego, CA).

## Graphics

Graphical assets made in © BioRender - biorender.com.

## Acknowledgments

We thank members of Hardy, Newton, Danthi, Mukhopadhyay, and Patton Labs for critical proofreading of the manuscript and for fostering ideas through helpful discussions. In addition, we thank Dr. David Mackenzie-Liu for generating the original cloning vector with the SacI restriction site. Finally, we thank our colleagues outside of IU for their generosity in sharing reagents, including Dr. Horacio Frydman, Boston University, for providing us Aa23 and C710 *Aedes albopictus* cells, Dr. William Sullivan for providing us with JW18 *Drosophila melanogaster* cells, and Dr. Zhiyong Xi, Michigan State University for providing us *Aedes aegypti* mosquitoes. This work was supported by NSF award to ILGN and RWH (MTM2025389), NIH R01 to ILGN (R01AI144430), NIH R21 to RWH (R21AI153785).

