## Supplementary material for "Differential viral RNA methylation contributes to pathogen blocking in *Wolbachia*-colonized arthropods": all supplemental material

Dataset S1 Legend: This excel spreadsheet contains raw data collected from the Incucyte S3 Live Cell Imaging Platform. The full dataset has been uploaded to dryad and can be accessed here.

| Primer Name | Forward Primer Sequence (5'-3') | Reverse Primer Sequence (5'-3') |
|---|---|---|
| CHIKV E2 | GGAATAAAGACGGATGATAGC | GGTCGGGAATGAAATTTTTCC |
| SINV E1 | TCAGATGCACCACTGGTCTCAACA | ATTGACCTTCGCGGTCGGATACAT |
| SINV nsP1 | AAGGATCTCCGGACCGTACTTG | CATGAACTGGGTGGTGTCGAAGC |
| Aedes 18S | CGAAAGTTAGAGGTTCGAAGGCGA | CCGTGTTGAGTCAAATTAAGCCGC |
| WSP | CATTGGTGTTGGTGTTGGTG | ACCGAAATAACGAGCTCCAG |
| Aedes GAPDH | CCGCTGATCTGCTAAACATAGA | GTTCTTCCGGGAGGATTCATTAG |
| Fly Mt2 | CCGTGGCGTGAAATAGCG | ACACCGCTTTCGGAGGACG |
| Aedes AMt2 | TATCAATCCGGTGGCCAATAC | CGGCGGTGACATGAGAATAA |
| pAFW-Mt2 *QC*_SalI | ACAAGGATGACGATGACAAGGTCCGAC | GGGTCGGCGCGCCCACCCTTGTCGAC |
| pAFW-Mt2_GA_Insert | AGGATGACGATGACAAGGTCATGGTATTTCGGGTCTTAGA | TCGGCGCGCCCACCCTTGTCTCATTTTATCGTCAGCAATT |
| pAFW-AMt2 | GCAACCGGTTTATGAGTGTTACCGACGGA | GCAGCTAGCTCAGTCCATCTCATCAAACAACGAACTC |
| Mt2-C78A_QC | GTCCCCGCCAGCTCAGCCCCACAC | ATCAGCAGCATGTTGGCC |
| AMt2-C78G_QC | GTCACCGGGCCAACCATTCA | ATGAGAGTAACGTTCACACCAAGCTTCTGAATG |

Table S1. Primers used in this study. Primers were purchased from Integrated DNA Technologies (IDT). All primers were used at a final concentration of 10μM for quantitative PCR and RT-PCR reactions. Recommended primer concentrations according to manufacturer's protocol were used for cloning experiments.

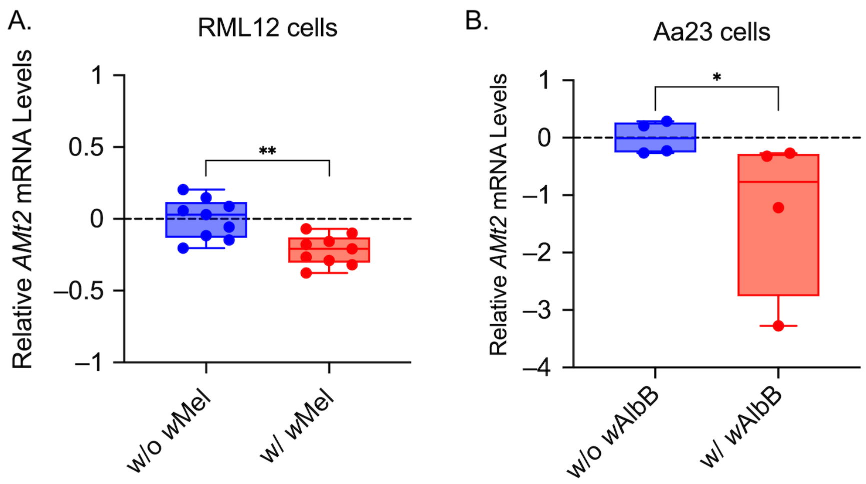


**Fig. S1. Presence of *Wolbachia* reduces MTase expression *ex vivo.*** Relative *AMt2* expression in the presence (w/ Wolb) and absence (w/o Wolb) of *Wolbachia* in *Aedes albopictus* cells. Quantitative RT-PCR was used to measure relative mRNA levels of *AMt2* in *Aedes albopictus* cells colonized with (A) *w*Mel strain of *Wolbachia* (RML12) and (B) *w*AlbB strain of *Wolbachia* (Aa23). Unpaired Mann Whitney U-tests on log-transformed values; RML12-*w*Mel $p = 0.0028$, Aa23-*w*AlbB $p = 0.0286$. Error bars represent standard error of mean (SEM) of independent experimental replicates. Primer details are available in Table S1. $^{**}P < 0.01$, $^{*}P<0.05$.

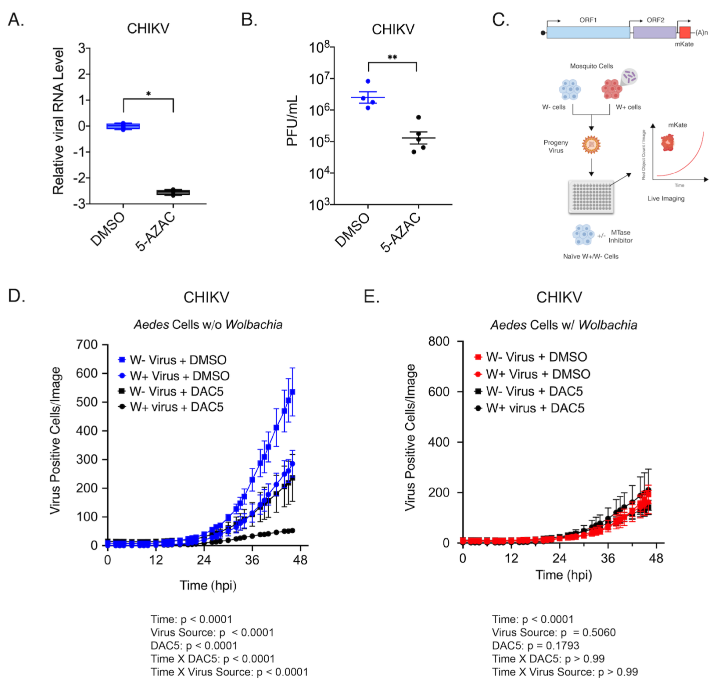


**Fig S2. Pharmacological inhibition of mosquito DNMT2 reduces virus replication and spread in mosquito cells.** Inhibition of mosquito DNMT2 in *Wolbachia*-free *Aedes albopictus* derived C710 cells was carried out using MTase inhibitors 5-Azacytidine (5-AZAC), or 5-Deoxyazacytidine (DAC5). Dimethyl-sulfoxide (DMSO) was used as the negative control. In each case, cells were pretreated with 5 μM inhibitors overnight prior to infections with CHIKV-mKate virus at MOI of 10. Cell lysates and supernatants were harvested at 48 hours post infection to quantify cellular viral RNA levels and infectious titer, respectively. (A) Levels of CHIKV RNA in mosquito cells treated with MTase inhibitor 5-AZAC were determined using quantitative RT-PCR. Unpaired two-tailed t-test with Welch's correction, CHIKV viral RNA: $p < 0.0001$, $t = 35.30$, $df = 6.001$. (B) Infectious CHIKV titers produced from mosquito cells treated with MTase inhibitor 5-AZAC were determined using plaque assays on BHK-21 cells. Unpaired two-tailed t-test with Welch's correction, CHIKV titer: $p = 0.0019$, $t = 4.864$ $df = 6.940$ (C) Schematic representation of live cell experiments. (D) CHIKV expressing mKate fluorescent reporter protein was grown in C710 *Aedes albopictus* cells in the presence (W+ virus) or absence (W- virus) of *Wolbachia (*strain *w*Stri). These progeny viruses were then used to infect naïve C710 cells without (D) and with (E) *Wolbachia (*strain *w*Stri) pretreated with MTase inhibitor (depicted in color) or DMSO (depicted in black) synchronously at a MOI of 1 particle/cell. Virus growth in cells was measured in real time by imaging and quantifying the number of red cells (Virus Positive Cells/Image) expressing the virus encoded mKate protein over a period of forty-eight hours, using live cell imaging. Color of the data points distinguish treatment conditions; blue represent C710 *Wolbachia*-free cells treated with DMSO, red represent C710 *Wolbachia*-colonized cells treated with DMSO, black represent both C710 cell types treated with 5 μM DAC5. Shape of data points represent the progeny virus type used to initiate infection; boxes represent viruses derived from W- cells, circles represent viruses derived from W+ cells. The Y-axis label red object count/Image represent virus-positive cells in a single field of view, four of which were collected and averaged/sample at each two-hour time point over the course of infection. Three-way ANOVA with Tukey's post hoc test for multivariate comparisons. Error bars represent standard error of mean (SEM) of independent experimental replicates (n=3). Three-way ANOVA with Tukey's multivariate analyses, DAC5: $p < 0.0001$, Virus Source: $p < 0.0001$, Time: $p < 0.0001$, DAC5 X Time: $p < 0.0001$, Source X Time: $p < 0.0001$, Virus Source X DAC5: $p = 0.1148$, Virus Source X DAC5 X Time: $p > 0.999$ (Fig S3D). Three-way ANOVA with Tukey's multivariate analyses, DAC5: $p = 0.1793$, Virus Source: $p = 0.5060$, Time: $p < 0.0001$, DAC5 X Time: $p > 0.99$, Virus Source X Time: $p > 0.99$, Virus Source X DAC5: $p = 0.1039$, Virus Source X DAC5 X Time: $p = 0.9804$ (Fig S3E). *P < 0.05; **P < 0.01, ****P < 0.0001.

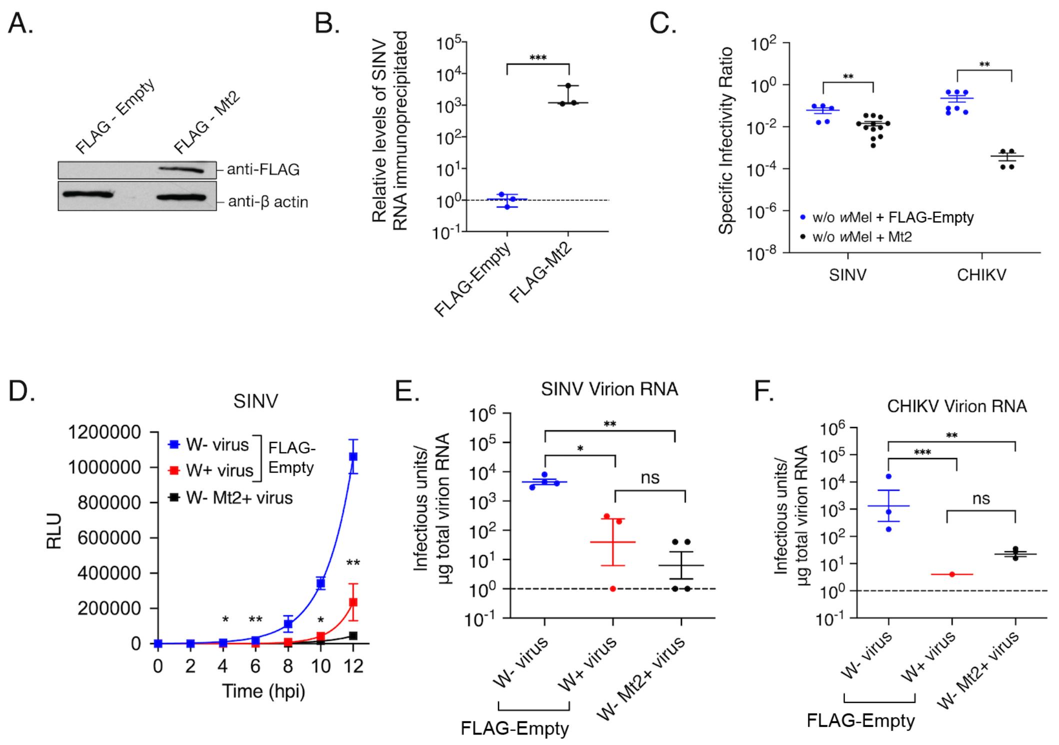


**Fig S3. *Drosophila melanogaster* DNMT2 functions as an antiviral.** (A) Western Blot of fly DNMT2 in *Wolbachia*-free *Drosophila melanogaster* derived JW18 cells transfected with expression vector constructs with (FLAG-Mt2) or without (FLAG-empty) *Mt2.* Cytoplasmic lysates of cells were collected 72 hours post transfection and probed with anti-FLAG and anti-β actin antibodies. (B) Relative levels of viral RNA recovered following AZA-IP of Mt2 in fly cells was quantified using qRT-PCR. JW18 fly cells without *Wolbachia* were transfected with expression vectors FLAG-empty or FLAG-Mt2 for 72 hours prior to infection with SINV at MOI of 10. Cells were treated for approximately 18h with 5 µM 5-Azacytidine to covalently trap Mt2 with its target cellular RNA prior to RNA immunoprecipitation using anti-FLAG antibody. One-sample two-tailed t-test performed on log-transformed values, $p = 0.001$, $t = 4.462$, $df = 11$. (C) Specific Infectivity Ratios of progeny viruses derived from *Drosophila melanogaster* cells colonized with native *Wolbachia* strain *w*Mel. Fly cells without *Wolbachia* were transfected with expression vectors FLAG-empty (w/o Wolb) or FLAG-Mt2 (w/o Wolb + Mt2) for 48 hours prior to infection with SINV-nLuc or CHIKV (MOI=10). Specific Infectivity Ratios of the progeny viruses generated 96 hours post infection were calculated as before. Unpaired two-tailed t-test with Welch's correction, SINV, $p = 0.0045$, $t = 3.698$, $df = 9.458$, CHIKV, $p < 0.0001$, $t = 9.608$, $df = 6.926$. (D) Progeny SINV-nLuc derived from fly cells with (W+ virus), without (W- virus) *Wolbachia* or overexpressing Mt2 (W- Mt2+ virus) were subsequently used to synchronously infect naïve BHK-21 cells at equivalent MOIs of 5 particles/cell. Cell lysates were collected at indicated times post infection and luciferase activity (RLU), was used as a proxy for viral replication. Two-way ANOVA Tukey's multiple comparisons tests, Time: $p < 0.0001$, *Wolbachia/AMt2*: $p = 0.0002$, Time X *Wolbachia/AMt2*: $p < 0.0001$. (E and F) Overexpression of *Drosophila* DNMT2 ortholog reduces infectivity of progeny virion RNA. Approximately 1µg of virion encapsidated RNA extracted from the aforementioned W+, W- and W- Mt2+ SINV (E) or CHIKV (F) viruses were transfected into naïve BHK-21 cells and infectious titer was determined by the counting the number of plaques produced after 48 hours post transfection. One-way ANOVA with Tukey's post hoc test for multivariate comparisons. Dotted line at Y=0 indicate the limit of detection. . Error bars represent standard error of mean (SEM) of independent experimental replicates. *P < 0.05; **P < 0.01, ***P < 0.001, ns = non-significant.

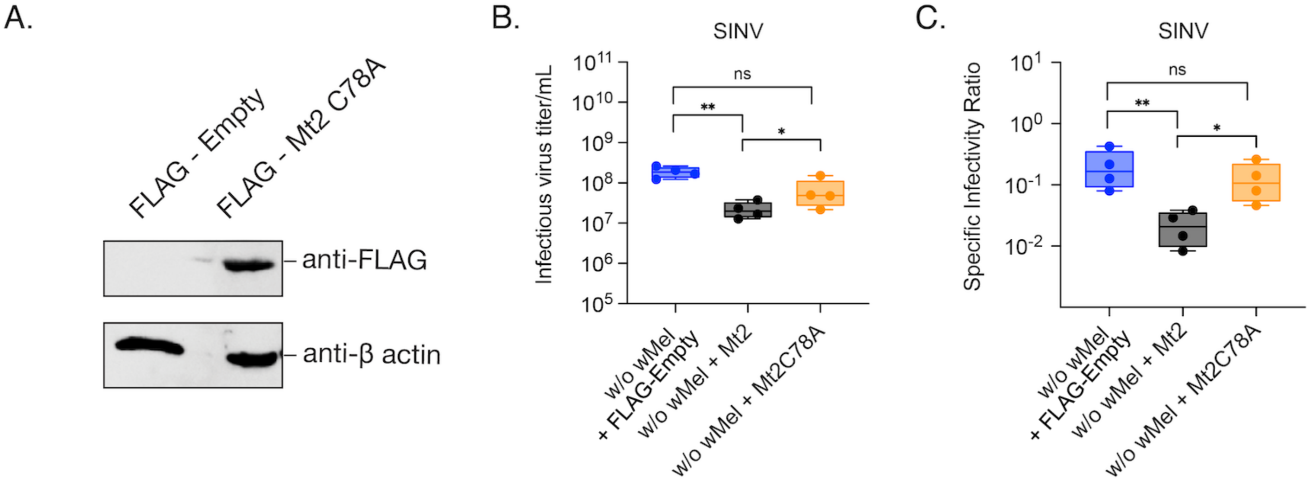


**Fig S4. Antiviral effect of fly MTase is dependent on its catalytic activity.** (A) Expression of the catalytic mutant of fly DNMT2 in *Wolbachia*-free *D. melanogaster* JW18 cells was assessed by Western Blot 72 hours post transfection with either the epitope tagged Mt2 mutant (FLAG-Mt2 C78A) or the empty control vector (FLAG-Empty) plasmid. (B) 72 hours after *Wolbachia*-free *D. melanogaster* JW18 cells were transfected with plasmids carrying either the wild-type (FLAG-Mt2), catalytic mutant (FLAG-Mt2 C78A) or the empty control vector (FLAG-Empty), they were challenged with SINV at MOI of 10. Cell supernatants were harvested 48 hours post infection, clarified, and used to assess Infectious SINV titer by standard plaque assay on vertebrate BHK-21 cells. One-way ANOVA with Tukey's post hoc test for multivariate comparisons. Error bars represent the standard error of mean of independent experimental replicates. (C) Specific Infectivity Ratios of progeny viruses produced 48 hours post infection was measured as the ratio of infectious virus titer (presented in B) to viral genome copies present in the cell supernatant, quantified using qRT-PCR using primers probing SINV E1 gene (see Materials and Methods for more details on the procedure and Table S1 for primer details). One-way ANOVA with Tukey's post hoc test for multivariate comparisons. Error bars represent the standard error of mean of independent experimental replicates. *P < 0.05, **P < 0.01, ns = non-significant.

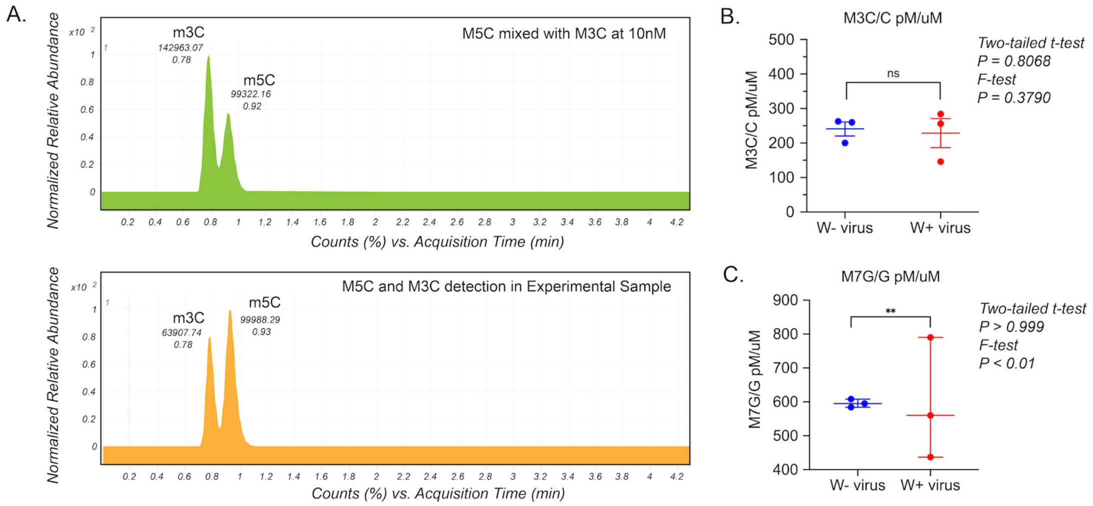


**Fig S5. LC-MS/MS detection of methylated cytosine residues.** (A) Chromatograms representing simultaneous detection of 3-methyl cytosine (m3C) and 5-methyl cytosine (m5C) groups in mixture comprised of 10nM of each standard (Top) and one representative experimental sample (Bottom). (B) Normalized 3-methyl cytosine content of RNA isolated from W- and W+ viruses represented as a ratio of total unmodified cytosine content. Unpaired two-tailed t-test, p = 0.8068, t = 0.2612, df = 4 (C) Normalized 7-methyl guanosine content of RNA isolated from W- and W+ viruses represented as a ratio of total unmodified guanosine content. Unpaired t-test with Welch's correction and F-test to compare variances. Error bars represent standard error of mean (SEM) of three independent virus preps from each cell type. F-test results: **P < 0.01, ns = non-significant.
